# Haploinsufficiency of A20 impairs protein-protein interactome and leads into caspase-8-dependent enhancement of NLRP3 inflammasome activation

**DOI:** 10.1101/401034

**Authors:** Kristiina Rajamäki, Salla Keskitalo, Mikko Seppänen, Outi Kuismin, Paula Vähäsalo, Luca Trotta, Antti Väänänen, Virpi Glumoff, Paula Keskitalo, Riitta Kaarteenaho, Airi Jartti, Nina Hautala, Päivi Jackson, Dan C Nordström, Janna Saarela, Timo Hautala, Kari K Eklund, Markku Varjosalo

## Abstract

**Objectives:** *TNFAIP3* encodes A20 that negatively regulates nuclear factor kappa light chain enhancer of activated B cells (NF-κB), the major transcription factor coordinating inflammatory gene expression. *TNFAIP3* polymorphisms have been linked with a spectrum of inflammatory and autoimmune diseases and recently, loss-of-function mutations in A20 were found to cause a novel inflammatory disease ‘haploinsufficiency of A20’ (HA20). Here we describe a family with HA20 caused by a novel *TNFAIP3* loss-of-function mutation and elucidate the upstream molecular mechanisms linking HA20 to dysregulation of NF-κB and the related inflammasome pathway.

**Methods:** NF-κB activation was studied in a mutation-expressing cell line using luciferase reporter assay. Physical and close-proximity protein-protein interactions of wild-type and *TNFAIP3* p.(Lys91*) mutant A20 were analyzed using mass spectrometry. NF-κB –dependent transcription, cytokine secretion, and inflammasome activation were compared in immune cells of the HA20 patients and control subjects.

**Results:** The protein-protein interactome of p.(Lys91*) mutant A20 was severely impaired, including interactions with proteins regulating NF-κB activation, DNA repair responses, and the NLR family pyrin domain containing 3 (NLRP3) inflammasome. The p.(Lys91*) mutant A20 failed to suppress NF-κB signaling, which led to increased NF-κB –dependent proinflammatory cytokine transcription. Functional experiments in the HA20 patients’ immune cells uncovered a novel caspase-8-dependent mechanism of NLRP3 inflammasome hyperresponsiveness that mediated the excessive secretion of interleukin-1β and -18.

**Conclusions:** The current findings significantly deepen our understanding of the molecular mechanisms underlying HA20 and other diseases associated with reduced A20 expression or function, paving the way for future therapeutic targeting of the pathway.

## INTRODUCTION

Tumor necrosis factor α induced protein 3 (TNAP3, also known as A20), encoded by the *TNFAIP3* gene, is a key negative regulator of NF-κB activation downstream a large number of immune receptors, including the TNF and Toll/interleukin(IL)-1 receptor families and the T and B cell antigen receptors^1-5^. A20 achieves this via its dual function as a ubiquitin-editing enzyme: an N-terminal ovarian tumor (OTU) domain removes activating K63-linked polyubiquitin from key intermediate NF-κB signaling molecules to destabilize protein-protein interactions, whereas a C-terminal zinc finger domain catalyzes the attachment of K48-linked polyubiquitin to induce proteasomal degradation^3 6 7^. In addition, A20 inhibits NF-κB activation via a noncatalytic mechanism involving its binding to linear ubiquitin on NF-κB essential modulator (NEMO)^8 9^. The mRNA expression of A20 is NF-κB –inducible^10^, thus generating a negative feedback loop to attenuate NF-κB responses.

Recently Zhou *et al.* reported six families with heterozygous loss-of-function mutations in the *TNFAIP3* gene that led to haploinsufficiency of A20 (HA20) and caused an early-onset Behçet-like disease^11^. Studies in mouse models have previously demonstrated a crucial role for A20 in auto-inflammation and autoimmunity, as mice with a germ-line deletion of *Tnfaip3* spontaneously develop severe multi-organ inflammation and tissue damage resulting in perinatal death^12^, and cell-specific ablation of *Tnfaip3* results in diverse symptoms of immune dysregulation^13 14^. Similar to *Tnfaip3* deficiency in mice^12-15^, human HA20 led to exaggerated NF-κB responses and increased secretion of NLR family pyrin domain containing 3 (NLRP3) inflammasome target cytokines^11^. Additional reports from these^16^ and further identified patients^17-21^ have expanded the clinical spectrum of HA20 to comprise diseases such as autoimmune lymphoproliferative syndrome, systemic juvenile arthritis, psoriatic arthritis, Crohn’s disease, and Hashimoto’s thyroiditis. The increasing number of diseases associated with A20 have emphasized its role in inflammatory diseases and in autoimmunity and raised the possibility that mutations in *TNFAIP3* may be an unexpectedly common underlying factor in the pathogenesis of rheumatic and other autoimmune diseases.

We describe here a family with polyautoimmunity and autoinflammatory symptoms caused by a novel heterozygous *TNFAIP3* p.(Lys91*) loss-of-function mutation resulting in haploinsufficiency. Our aim was to further elucidate the upstream molecular mechanisms linking HA20 to dysregulated NF-κB activation and NLRP3 inflammasome responses. Thus, we analyzed protein-protein interactions of wild-type (wt) and p.(Lys91*) A20 in HEK cells and performed functional experiments in patients’ immune cells to study NLRP3 inflammasome responses. As A20 is rapidly emerging as a key regulator of immune activation and autoimmunity in humans, the detailed characterization of A20 effector mechanisms paves the way for future therapeutic targeting of the pathway.

## MATERIALS AND METHODS

### Ethics statement

Blood samples from patients and controls were collected under written consent in accordance with the Declaration of Helsinki; also the parents of the pediatric subjects signed a written informed consent. The study protocol was approved by the Ethical Committee for Clinical Science of Oulu University Hospital and by the Coordinating Ethics Committee of The Hospital District of Helsinki and Uusimaa.

### Genetic analyses

DNA samples were extracted from total peripheral blood using standard methods. Whole-exome sequencing and data analysis were performed as previously described.^22 23^ All the common (frequencies above 0.01 in the general population) and non-coding variants were discarded. We searched for rare heterozygous variants, according to the inferred autosomal dominant inheritance pattern (**Fig. 1A**). The variant identified in the *TNFAIP3* gene (Ensembl ENSG00000118503:ENST00000433680: exon2:c.A271T: p.(Lys91*) was confirmed using PCR and capillary electrophoresis.

**Figure 1.**
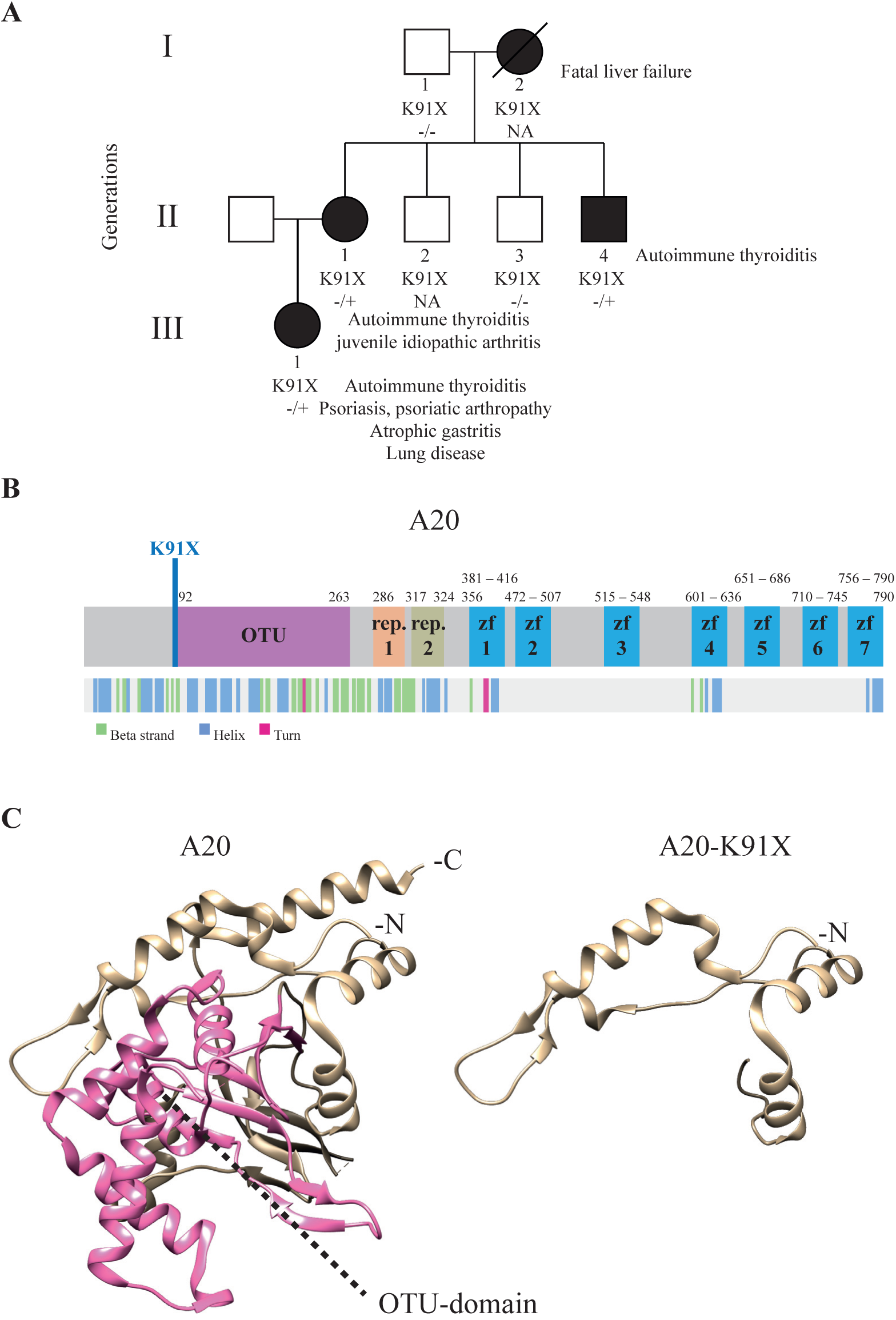
Phenotypic and protein level characteristics of a family with newly identified *TNFAIP3* c.A271T: p.(Lys91*) mutation. A) A pedigree showing the *TNFAIP3* p.(Lys91*) mutation carriers (-/+, solid symbols) in three generations (I-III). Autoimmune thyreoiditis was diagnosed in all carriers (I-2, II-1, II-4, III-1) at an early age. In addition, the index patient (II-1) suffered from psoriasis, articular symptoms, atrophic gastritis, severe autoinflammatory lung reaction, anemia, genital papillomatosis, and repeated genital HSV infections. Daughter of the index (III-1) was diagnosed with polyarticular juvenile idiopathic arthritis at the age of 4 years, and mother of the index (I-2) died of liver failure at the age of 46 years. B) Schematic illustration of A20 structure showing the protein domains (upper panel) and the folding (lower panel). Position of the novel p.(Lys91*) mutation is indicated in blue. C) 3D structure models of wild-type and mutant p.(Lys91*) A20 (based on PDB accession 5LRX). OTU domain is highlighted with pink.

### Creating A20 expressing Flp-In 293 T-REx cell lines

*A20* mutant plasmids were created using Quick-Change Site-Directed Mutagenesis kit (Agilent Technologies) to wild-type *TNFAIP3* obtained from the human Orfeome collection (Horizon Discovery). TNFAIP3 constructs were further subcloned into N-terminal MAC-tag Gateway® destination vector^24^. All generated constructs were confirmed with direct sequencing. The MAC-tagged expression constructs were transfected into Flp-In 293 T-REx cells (Invitrogen, Life Technologies) with FuGENE® HD (Promega). The cells were grown according to manufacturer’s instructions under selection with Hygromycin B (Thermo Fisher Scientific) to create stable cell lines. Expression of stable transgenes was confirmed by Western blotting using anti-hemagglutin (HA) primary (Biolegend, MMS-101R) and horseradish peroxidase-conjugated secondary antibody.

### Luciferase assay

For reporter assays HEK293 cells were transfected with in total 50 ng A20 mutant or wild-type MAC-tagged construct and GFP or empty MAC-tag-vector^24^, 40 ng of NF-κB luciferase reporter plasmid (Cignal 45), and 1.4 ng Renilla luciferase reporter plasmid (pRL-SV40). Transfected cells were stimulated with indicated amounts of TNF-α (R&D Systems) for 16 hours, lysed with 1x passive lysis buffer (Promega), and luciferase assays performed with Dual-Glo Luciferase Assay System according to manufacturer’s instructions (Promega).

### Affinity Purification and mass spectrometry

For each pull-down approximately 5×10^7^ cells (5 × 15 cm dishes) were induced with 2μg/ml tetracycline (Thermo Fisher Scientific) for 24 hr (for BioID additional 50 μM biotin was added). After induction, cells were washed, harvested, pelleted by centrifugation, snap-frozen, and stored at -80 °C until analysis. Affinity purification of AP-MS and BioID was performed as described in *Turunen et al.* 25 and *Heikkinen et al.* ^26^, respectively. Sample preparation for mass spectrometry and LC-MS/MS analysis on Orbitrap Elite ETD mass spectrometer was performed as in *Heikkinen et al.* ^26^. After peptide reconstitution, 4 μl were analyzed from both Strep-Tag and BioID-samples.

### Culture and stimulation of peripheral blood mononuclear cells (PBMCs)

PBMCs were isolated by density gradient centrifugation in Ficoll-Paque PLUS (GE Healthcare) and seeded at 1.5 × 10^6^ cells/ml in Macrophage-SFM medium supplemented with penicillin-streptomycin or in RPMI 1640 supplemented with GlutaMAX, penicillin-streptomycin, and 10 % FBS (all from Gibco). PBMCs were allowed to rest for a minimum of 3 h before starting the stimulations with 1 μg/ml lipopolysaccharides (LPS) from *E.coli* O111:B4 (Sigma), 1 μg/ml Pam_3_Cys-SKKKK (Pam-_3_Cys; EMC microcollections), 10 μg/ml polyinosinic-polycytidylic acid (poly(I:C); InvivoGen), and 5 mM adenosine 5’-triphosphate (ATP; Sigma) for the indicated times. Inhibitors Z-YVAD-FMK (20 μM; R&D Systems), Z-IETD-FMK (5 μM; BD Bioscience), and necrostatin-1 (30 μM; Enzo Life Sciences) were added to the cells simultaneously with LPS in the 16 h stimulations, or 30 min before LPS in the LPS 6 h +ATP 45 min stimulations.

### Measurement of cytokine secretion

TNF-α and the mature, cleaved forms of IL-1β and IL-18 were detected from PBMC culture media supernatants using Human TNF-α DuoSet ELISA, Human IL-1β/IL-1F2 DuoSet ELISA, and Human Total IL-18 DuoSet ELISA (all from R&D Systems).

### Quantitative real-time PCR

RNA was isolated from PBMCs using RNeasy Plus Mini Kit (Qiagen) and cDNA synthesized with iScript kit (Bio-Rad). Quantitative real-time PCR was performed from 10 ng of cDNA per reaction using LightCycler480 SYBR Green I master (Roche) and LightCycler96 instrument (Roche); primer sequences are listed in **Supplementary table 1**. Relative gene expression was calculated using the 2^(-δδCt)^method, normalizing to the geometric mean of expression of two housekeeping genes, ribosomal protein lateral stalk subunit P0 and beta-2 microglobulin.

### Monocyte caspase-1 activity measurement

Aliquots of EDTA-anticoagulated blood were incubated for 6 h at +37 °C under gentle rocking with or without 10 ng/ml LPS (Sigma). ATP at 5 mM (Sigma) was added for 20 min, followed by 2h incubation at +37 °C with a cell-permeable, irreversibly binding FAM-FLICA fluorescent caspase-1 substrate (ImmunoChemistry Technologies). After red blood cell lysis and staining with the monocyte marker anti-human CD14-APC (Miltenyi Biotec) for 15 min at +4 °C, the cells were analyzed using BD Accuri C6 flow cytometer (BD Biosciences; instrument maintained by the Biomedicum Flow Cytometry Unit).

## RESULTS

### Identification of the mutation and the clinical manifestations of the patients

Exome sequencing of the index patient (II-1) identified a novel nonsense mutation in the exon 2 of the *TNFAIP3* gene: c.A271T: p.(Lys91*) (ENSG0 0000118503:ENST00000433680)(**Fig. 1A-B**). The mutation is not listed in Genome Aggregation Database (gnomAD, Cambridge, MA, USA), and was predicted to be pathogenic according to the ACMG Standards and Guidelines^27^. The *TNFAIP3* p.(Lys91*) mutation was confirmed in the patient’s genomic DNA and screened in the other members of the family. Two other affected family members (II-4, III-1) were observed to also harbor the mutation, (I-2 is an obligate carrier). The clinical manifestations of the carriers included autoimmune thyroiditis, which was diagnosed in all carriers (I-2, II-1, II-4, III-1) at an early age. The index patient suffered from psoriasis, articular symptoms, atrophic gastritis, severe inflammatory lung reaction, anemia and repeated genital HSV infections. Daughter of the index was diagnosed with polyarticular juvenile idiopathic arthritis at the age of 4. See also the legend of **Fig. 1A**.

### Effect of the TNFAIP3 p.(Lys91*) mutation on protein structure and expression

The 3D protein structures of wild-type (wt) and p.(Lys91*) A20 were visualized by processing the PDB (www.rcsb.org^28^) accession number 5LRX (chain A^29^) with Chimera^30^, which yielded a severely truncated protein lacking the complete OTU domain (**Fig. 1C**). To further analyze the expression and possible physical and functional changes caused by the *TNFAIP3* p.(Lys91*) mutation, stable Flp-In 293 T-REx cell lines expressing wt and p.(Lys91*) A20 were created. The wt HA-tagged A20 was expressed in Western blot, but the p.(Lys91*) mutant was detected only with an N-terminal tag (**Fig. 2A**), suggesting that only an N-terminal fragment is produced and no alternative start codons are used. Peptide identification using mass spectrometry confirmed that only the N-terminal region was detectable in the p.(Lys91*) mutant (**Fig. 2B**). The expression levels of the N-terminal peptides/protein of the wt and p.(Lys91*) mutant A20 were highly comparable (**Fig. 2B**).

**Figure 2.**
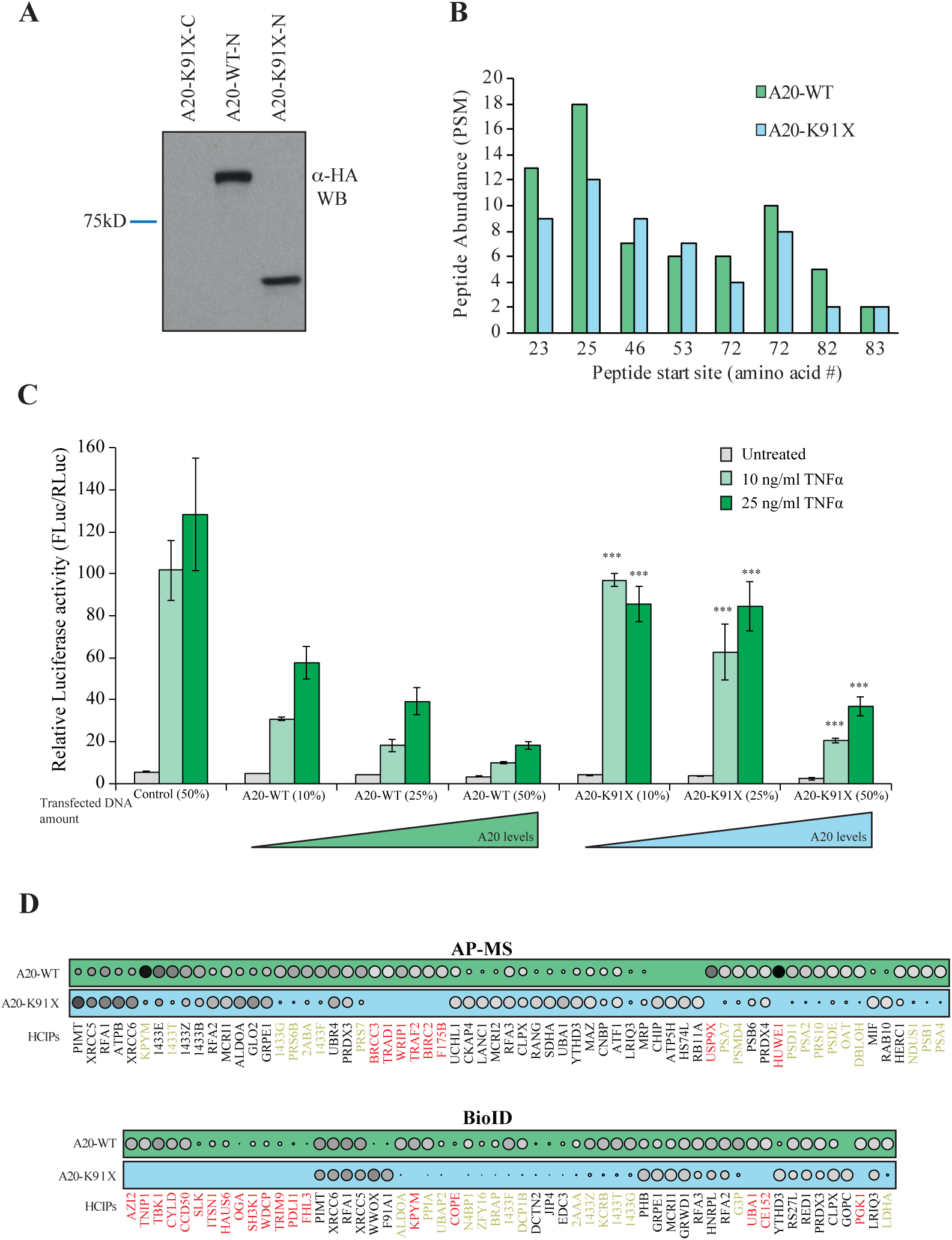
The p.(Lys91*) mutant A20 fragment is expressed but shows severe impairment of protein-protein interactions and of inhibitory activity on NF-κB signaling. A) Expression profile analysis of wt A20 and the p.(Lys91*) mutant in HEK293 cells. The wt protein is detected with both as N- and C-terminally –tagged constructs, whereas the p.(Lys91*) mutant is only detected as N-terminally –tagged illustrating that no alternative start site is used and only short N-terminal fragment is produced. B) Quantitative mass spectrometry-based proteomics analysis shows that the N-terminal part of the wt A20 and p.(Lys91*) mutant are expressed in similar levels as shown by the quantitative abundance of the peptides derived from the N-termini. C) wt A20 dose-dependently inhibits the NF-κB signaling induced by TNF-α (Key:light green=10ng/ml and dark green=25ng/ml), whereas the p.(Lys91*) mutant show clearly diminished inhibitory activity suggesting haploinsufficiency. The asterisks represent the significance values; * <0.05, ** <0.01, *** <0.001. D) AP-MS (upper panel) and BioID (lower panel) analysis of wt and p.(Lys91*) mutant A20 shows clear differences in their stable protein-protein interactions and BioID analysis displays differential molecular context of the p.(Lys91*) mutant in comparison with the wt A20. (Key: the size and the color gradient show the relative abundance of the interacting prey protein detected in association with A20, red typeface designates the complete loss of the corresponding interactions and yellow >50% decrease compared to the wt A20).

### A20 p.(Lys91*) mutant fails to suppress NF-κB activation and shows severely impaired protein-protein interactome

To analyze the possible functional consequences of the p.(Lys91*) mutation and the loss of OTU domain on the A20 functions, we monitored the NF-κB pathway activity using a luciferase-based reporter assay. The wt A20 dose-dependently inhibited TNF-α induced NF-κB pathway activation, whereas the p.(Lys91*) mutant failed to do so, clearly suggesting a haploinsufficiency (**Fig. 2C**).

The physical effects of the p.(Lys91*) mutation were further analyzed by proteome-wide affinity purification mass spectrometry (AP-MS) (see **Supplementary Fig. 1** for details). These analyses revealed that the p.(Lys91*) mutant completely loses physical interactions with BRCC3, TRAD1, WRIP1, TRAF2, BIRC2, F175B, USP9X and HUWE1, with several additional weakened interactions (**Fig. 2D**). To obtain more insight into the molecular mechanisms of A20 and especially on the p.(Lys91*) mutation, we further quantitatively analyzed the functional and close-proximity interactions using BioID MS^24^(see **Supplementary Fig. 1** for details). This analysis revealed complete loss of interactions with AZI2, TNIP1, TBK1, CYLD, CCD50, SLK, ITSN1, HAUS6, OGA, SH3K1, WDCP, TRIM9, PDLI1, FHL3, KPYM, COPE, UBA1, CE152 and PGK1. Many more interactions were severely diminished (**Fig. 2D**). Majority of the detected interactions of wt A20 were novel (**Fig. 3**). Functional annotation revealed extensive interactions with ubiquitinylation-, proteasome- and NF-κB -related proteins, as well as with proteins involved e.g. in cell fate decisions (14-3-3 proteins), DNA repair (53BP1 complex), and glycolysis (**Fig. 3**). Of these, the interactions with ubiquitinylation- and NF-κB -related proteins were most severely affected in the A20 p.(Lys91*) mutant.

**Figure 3.**
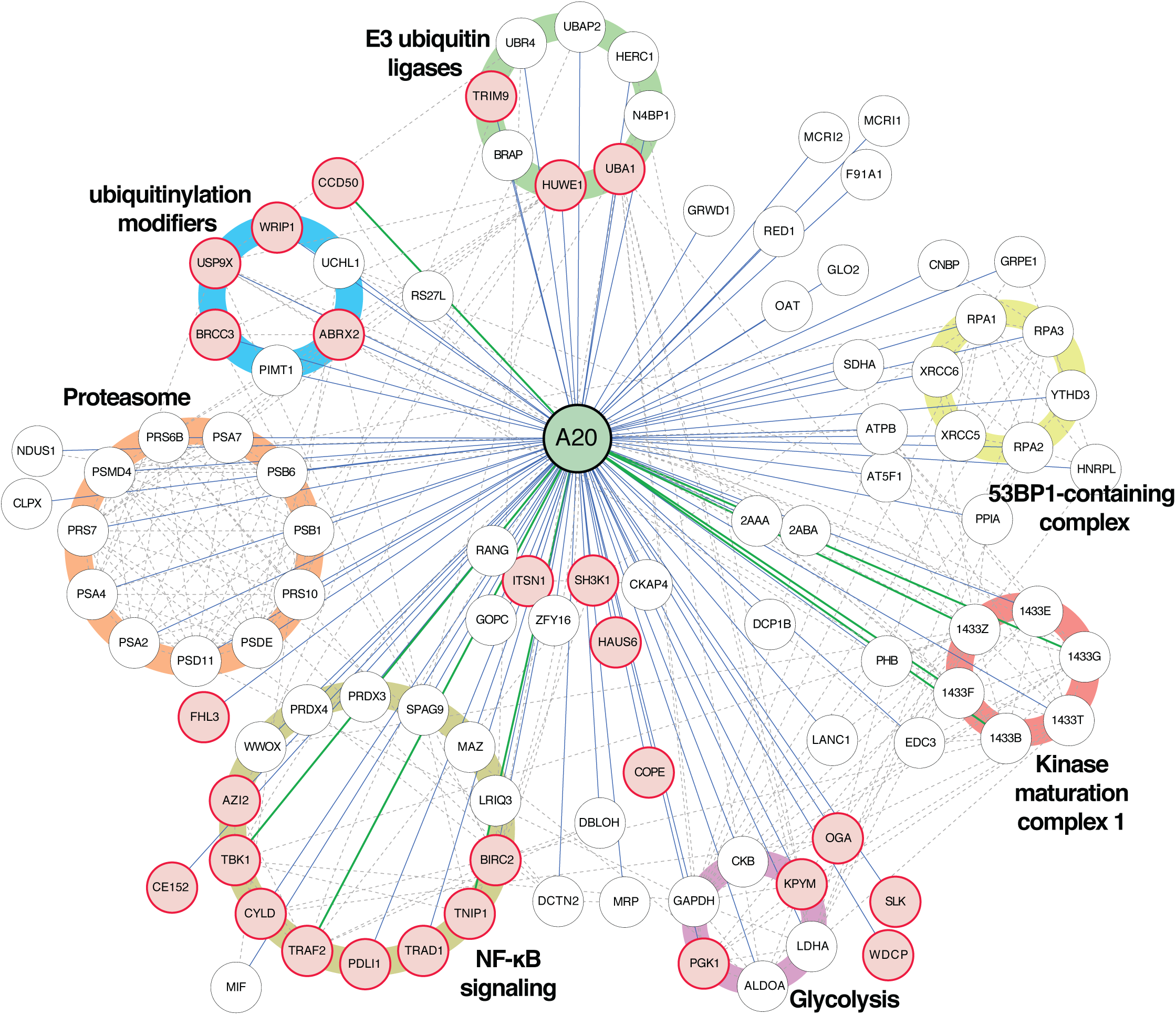
Interactome analysis reveals known and novel interactions for A20. Affinity purification mass spectrometry and BioID analysis of A20 identified 98 high-confidence protein-protein interactions (known interactions are shown with green lines, novel interaction detected on this study with blue and known prey-prey interactions with a dashed line). The interacting proteins are grouped bases on their molecular functions/complexes. Interactions, which are lost with the p.(Lys91*) mutation, are illustrated with red node fill color.

### Blood immune cells of TNFAIP3 p.(Lys91*) carriers display strongly increased secretion of inflammasome-controlled cytokines IL-1β and IL-18

The NF-κB pathway and A20 have been shown to play regulatory roles in the activation of NLRP3 inflammasome, a pathway triggering caspase-1-mediated proteolytic maturation and secretion of proinflammatory cytokines IL-1β and IL-18^15 31 32^. Excessive NLRP3-dependent cytokine secretion was demonstrated in HA20^11^, yet the mechanism(s) linking reduced A20 to NLRP3 inflammasome activation in patient cells were not elucidated. We stimulated peripheral blood mononuclear cells (PBMCs) of the HA20 patients with Toll-like receptor (TLR) 4 agonist LPS, with or without a subsequent ATP pulse; both stimuli trigger the NLRP3 inflammasome in monocytes via distinct mechanisms and with different kinetics^31^. We found elevated secretion of IL-1β and IL-18 in PBMCs of the *TNFAIP3* p.(Lys91*) carriers in response to LPS+ATP-induced ‘canonical’ (**Fig. 4A-B; left panels**) as well as LPS-induced ‘alternative’ pathway of NLRP3 inflammasome activation (**Fig. 4A-B; right panels**). To control for potential confounding factors in FBS preparations, PBMC cytokine secretion was analyzed both in commercial serum-free monocyte-macrophage medium (**Fig. 4**) and in RPMI containing 10 % FBS (**Supplementary Fig. 2**), yielding similar results. Also, TNF-α secretion in response to LPS, Pam_3_Cys, and poly(I:C) was increased in the patients (**Fig. 4C, Supplementary Fig. 2C**). We further studied the levels of IL-1β and IL-18 in patients’ plasma by ELISA and found increased levels of circulating IL-18 in *TNFAIP3* p.(Lys91*) carriers compared to the age- and sex-matched controls (**Supplementary Fig. 3A**), whereas IL-1β was not detectable.

**Figure 4.**
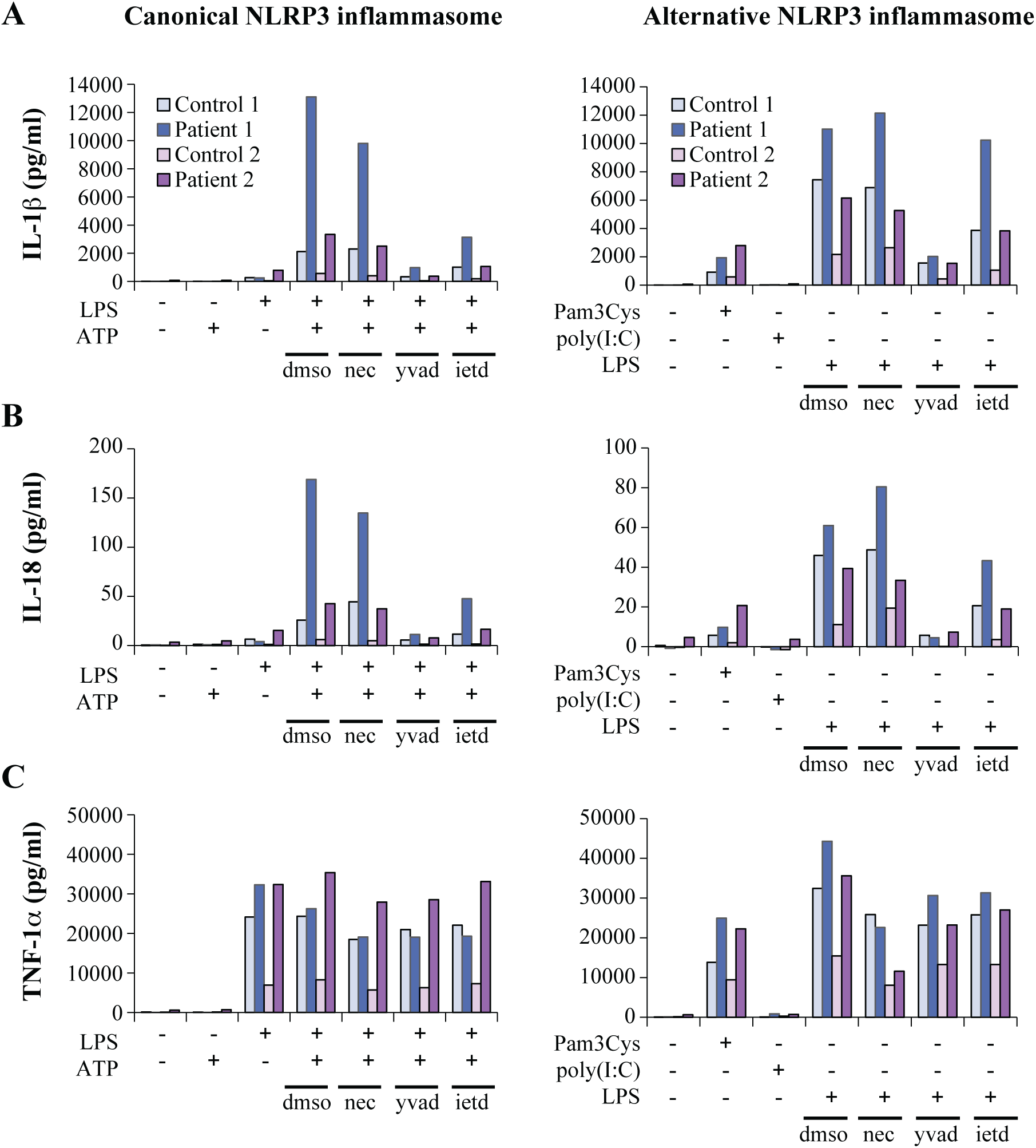
Inflammasome-dependent and –independent proinflammatory cytokine secretion is elevated in PBMCs of *TNFAIP3* p.(Lys91*) carriers. PBMCs from *TNFAIP3* p.(Lys91*) mutation carriers, patients 1 (II-1, female, 30 years old) and 2 (III-1, female, 8 years old), were compared to sex- and age-matched controls 1 and 2, respectively. (A-C, left panels) The cells were primed with LPS for 6 h, followed by treatment with ATP for 45 min. (A-C, right panels) The cells were treated with TLR agonists for 16 h. Inhibitors necrostatin-1 (nec), Z-YVAD-FMK (yvad), and Z-IETD-FMK (ietd), or solvent control dimethyl sulfoxide (dmso), were added as indicated and cytokines were detected from PBMC culture supernatants by ELISAs.

### The increased NLRP3 inflammasome response in TNFAIP3 p.(Lys91*) carriers is dependent on enhanced pro-IL-1β transcription and caspase-8 activity, but not on RIPK1

To elucidate the mechanism of altered NLRP3 inflammasome activation in the HA20 patients’ PBMCs, we analyzed the mRNA expression of inflammasome pathway components and cytokines and the effect of inflammasome-related inhibitors on cytokine secretion. PBMCs of patient III-1 displayed strongly increased baseline expression of pro-IL-1β and NLRP3 receptor, whereas patient II-1 cells showed only moderate changes (**Supplementary Fig. 3B**). After LPS stimulation, PBMCs from both patients showed slightly elevated levels of pro-IL-1β mRNA, and NLRP3 receptor expression remained moderately elevated only in patient III-1 (**Fig. 5A**). LPS-induced expression of NF-κB target cytokines IL-6, TNF-α, and IL-8 was elevated in patient III-1, and IL-6 also in patient II-1 (**Fig. 5B**). Receptor interacting serine/threonine-protein kinase 1 (RIPK1) inhibitor necrostatin-1 did not affect IL-1β or IL-18 secretion in the patients’ PBMCs (**Fig. 4A-B**), although it normalized the excessive TNF-α response to 16 h LPS treatment (**Fig. 4C, right panel**). As expected, both canonical and alternative NLRP3 inflammasome responses were efficiently blocked by the caspase-1 inhibitor Z-YVAD-FMK (**Fig. 4A-B**). However, levels of active caspase-1 in monocytes after whole blood inflammasome stimulation were increased only in patient II-1 (**Fig. 5C**), supporting a more indirect enhancement of inflammasome complex function. Remarkably, caspase-8 inhibitor Z-IETD-FMK efficiently suppressed IL-1β and IL-18 secretion in PBMCs of the *TNFAIP3* p.(Lys91*)- positive patients, particularly after canonical NLRP3 inflammasome activation (**Fig. 4A-B**).

**Figure 5.**
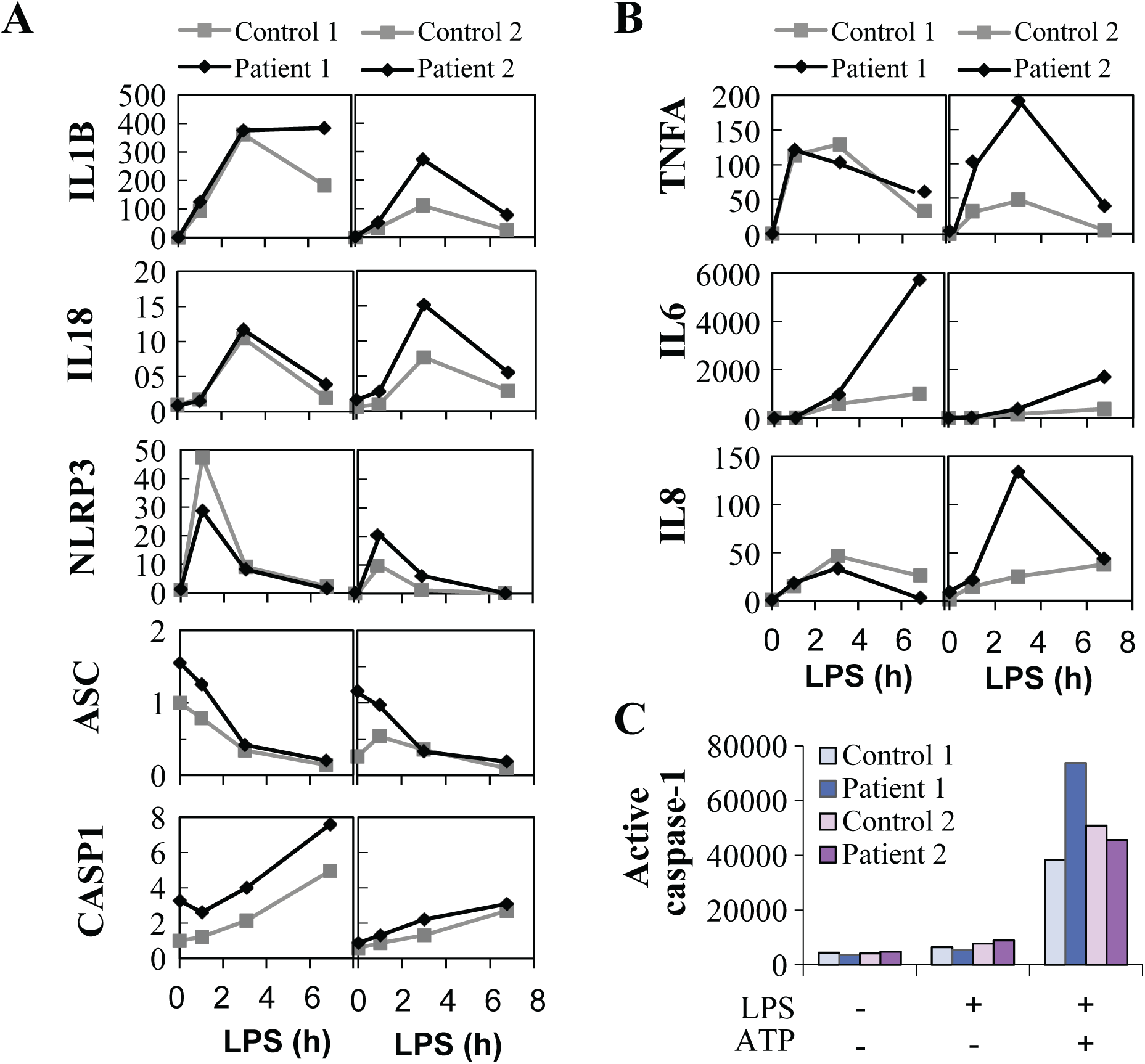
Expression and activation of the NLRP3 inflammasome is altered in PBMCs of *TNFAIP3* p.(Lys91*) carriers. Samples from *TNFAIP3* p.(Lys91*) mutation carriers, patients 1 (II-1, female, 30 years old) and 2 (III-1, female, 8 years old), were compared to sex- and age-matched controls 1 and 2, respectively. Relative expression of (A) NLRP3 inflammasome components and target cytokines and (B) inflammasome-independent proinflammatory cytokines was analyzed in PBMCs by quantitative PCR and normalized against housekeeping gene expression (arbitrary units). (C) Whole blood was stimulated with LPS and ATP, followed by the detection of NLRP3 inflammasome-triggered caspase-1 activity in monocytes using a fluorescent FLICA probe; the data are presented as median fluorescence intensity (MFI).

## DISCUSSION

HA20 is a recently described familial autoinflammatory disease originally reported to manifest as an early-onset Behçet-like disease^11^, yet the range of known clinical manifestations is rapidly expanding^16-21^. Here we studied patients with HA20 caused by a novel heterozygous *TNFAIP3* p.(Lys91*) mutation resulting in severe truncation and production of an N-terminal protein fragment with highly limited functional capacity. The patients exhibit an unusual combination of early onset autoimmune diseases, autoinflammatory symptoms and immunodeficiency, reflecting the widespread functions of A20 in controlling inflammation. To gain further insight into the molecular mechanism underlying the development of this complex disease, we focused on scrutinizing two pathways strongly linked with A20 function; the NF-κB pathway and the NLRP3 inflammasome.

Mutations causing HA20 were previously reported to increase TNF-α –induced IKKα/ IKKβ and MAPK phosphorylation and activation of NF-κB via defective removal of K63-linked ubiquitin from RIPK1, TNF receptor-associated factor 6, and NEMO^11 18 19^. As expected, also the A20 p.(Lys91*) mutant was defective in suppressing TNF-α –induced NF-κB activation. We employed state-of-the-art proteomics tools to comprehensively map global changes in A20 protein-protein interactions caused by the p.(Lys91*) mutation. In agreement with A20 ubiquitin-editing functions, the results revealed loss of physical interactions with ubiquitin ligases (BIRC2, TRAF2, HUWE1), deubiquitinases (BRCC3, USP9X), and other ubiquitinylation modulators (WRIP1, TRAD1, F175B/ ABRX2). BIRC2/3 and TRAF2/3 form a regulatory complex that suppresses apoptotic functions of caspase-8 to enforce canonical (inflammatory) NF-κB signaling, while blocking noncanonical NF-κB activation^33 34^. A20 modulates this balance by inducing TRAF2 degradation to suppress canonical NF-κB activation, and by binding c-IAP1/2 to support their caspase-8-inhibiting function while blocking their noncanonical NF-κB-inhibiting function^35-37^. Thus, the loss of A20 p.(Lys91*) interaction with BIRC2 and TRAF2 is compatible with enhancement of caspase-8 activity and canonical NF-κB signaling. In turn, BRCC3, F175B/ABRX2, WRIP1, and HUWE1 have specific functions in DNA damage responses^38-41^, which could be a significant finding regarding the association of reduced A20 levels with systemic lupus characterized by autoantibodies against nuclear antigens^14^. Moreover, BRCC3 and F175B/ ABRX2 are also components of the BRISC deubiquitinase complex that cleaves K63-linked ubiquitin and regulates mitotic spindle assembly^42^, type I interferon signaling^43^, and NLRP3 inflammasome activation^44^. Also, functional close-proximity protein-protein interactions of the A20 p.(Lys91*) mutant, particularly those related to ubiquitinylation and NF-κB signaling, were severely impaired, whereas interactions with proteasome, 53BP1-containing DNA repair complex, and 14-3-3 regulatory proteins remained partially functional.

The roles of A20 in regulation of the NLRP3 inflammasome, both NF-κB –dependent and –independent, are just beginning to be uncovered^15 32^. Mice deficient in myeloid A20 develop erosive polyarthritis specifically due to poor control of NLRP3 inflammasome activation by A20^15^. Moreover, increased NLRP3-dependent secretion of IL-1β and IL-18 was reported in HA20 patients^11 17^, yet the upstream mechanism(s) linking this effect to A20 in patient immune cells remain unknown. We found increased IL-1β and IL-18 secretion in PBMCs of *TNFAIP3* p.(Lys91*)- positive patients in response to both canonical and alternative NLRP3-activating stimuli. In mouse macrophages, A20 suppresses the NF-κB-induced mRNA expression of NLRP3 receptor and pro-IL- 1β, but also blocks RIPK1/3-dependent pro-IL-1β ubiquitination that supports the inflammasome-mediated processing into mature IL-1β^15 32^. Our patients’ PBMCs showed elevated LPS-induced transcription of pro-IL-1β, but NLRP3 receptor expression was not consistently increased. The RIPK1 inhibitor necrostatin-1 that reduced aberrant IL-1β secretion in *Tnfaip3-*deficient mouse macrophages^32^ had no effect on IL-1β nor IL-18 secretion in the patients’ PBMCs, which was, instead, strongly suppressed by caspase-8 inhibition. This suggests a novel caspase-8-dependent mechanism linking reduced A20 function to enhanced NLRP3 inflammasome activation. The finding is in line with the increased caspase-8 activity reported in *Tnfaip3*-deficient mouse cells^32 45^and with the caspase-8-suppressing functions of A20^37 45^, and further supported by the loss of physical interaction between A20 p.(Lys91*) mutant and BIRC2, which is required for caspase-8 inhibition by A20^37^. Previous studies suggest that alongside its apoptotic functions, caspase-8 facilitates the priming and activation of NLRP3 inflammasome via multiple mechanisms^46^. In addition, the loss of interaction between A20 p.(Lys91*) mutant and the BRCC3-containing BRISC complex suggests a further novel NLRP3- regulating function of A20 that is lost in haplo-insufficient cells, as BRCC3 deubiquitinates the NLRP3 receptor directly promoting its activation^44^.

In summary, we report here a *TNFAIP3* p.(Lys91*) loss-of-function mutation leading to HA20 and autoimmune/inflammatory symptoms. We utilize this mutation as a model to uncover novel molecular mechanisms related to the role of A20 in disease development. We defined, for the first time, the global physical and functional interactome of wild-type A20 and found severely diminished interactions in the p.(Lys91*) mutant. Moreover, we discovered a novel caspase-8- dependent mechanism linking HA20 to hyperactivation of the NLRP3 inflammasome in patient immune cells. Collectively, these data greatly deepen our understanding of the molecular mechanisms underlying the clinical disease phenotype of HA20 and other diseases associated with reduced A20 expression or function.

## ACKNOWLEDGEMENTS

The study was supported by Finska Läkaresällskapet (K.K.E., D.N), The Canadian Institutes of Health Research (THC 135230; K.K.E.), the Stockmann foundation (K.K.E.), and the Paulo foundation (K.R.). Molecular graphics and analyses were performed with the UCSF Chimera package. Chimera is developed by the Resource for Bio-computing, Visualization, and Informatics at the University of California, San Francisco (supported by NIGMS P41-GM103311). Exome sequencing of the index patient was conducted in the Institute for Molecular Medicine Finland (FIMM) Technology Centre. Sini Miettinen is thanked for expert mass spectrometry skills, and Xiaonan Liu for help with Chimera.

---

**Figure S1.**
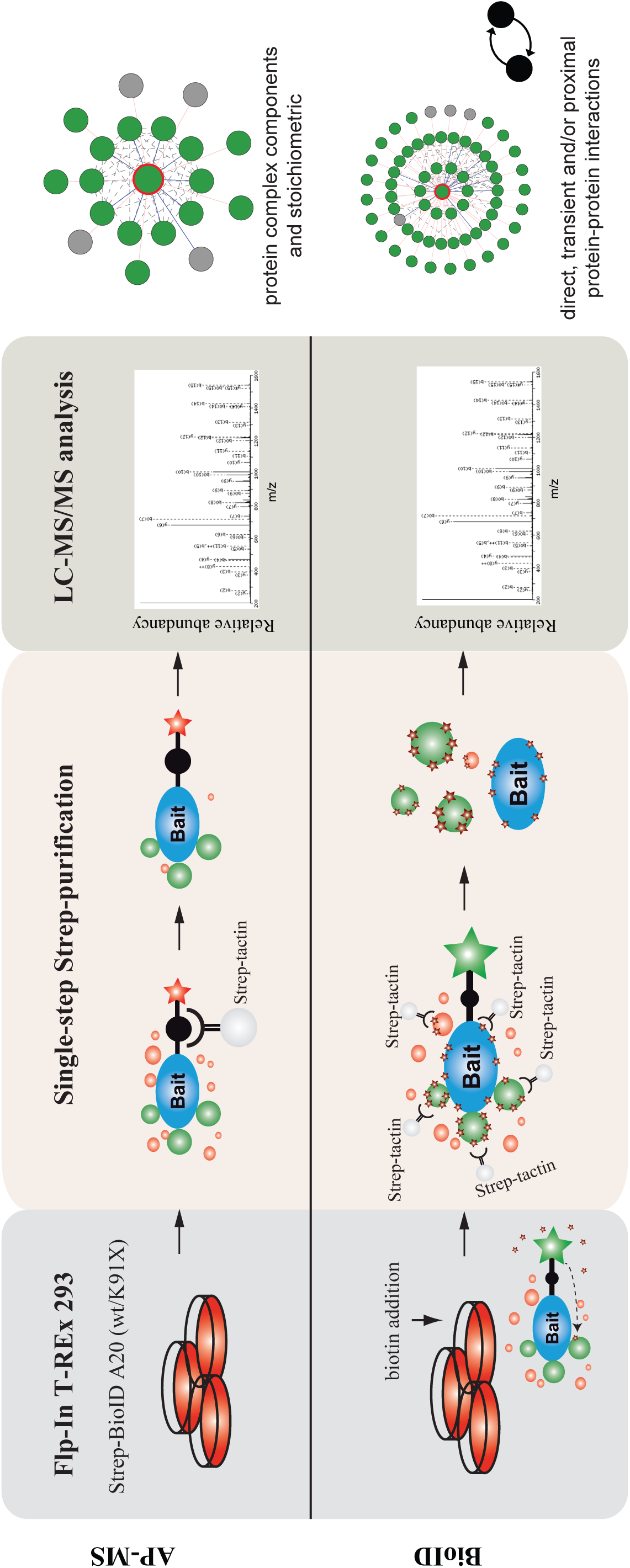
SUPPLEMENTARY FIGURE 1.

**Figure S2.**
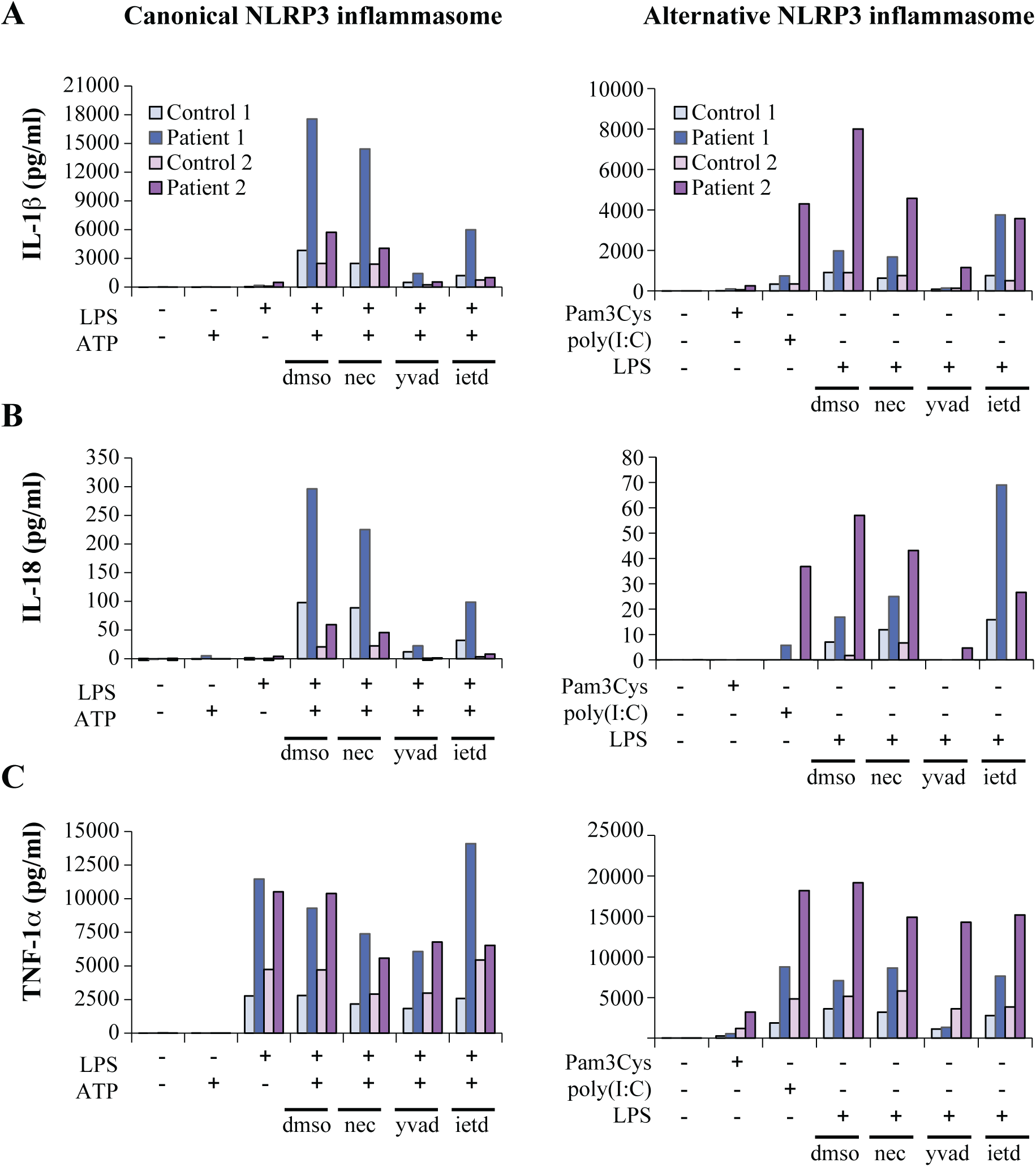
SUPPLEMENTARY FIGURE 2.

**Figure S3.**
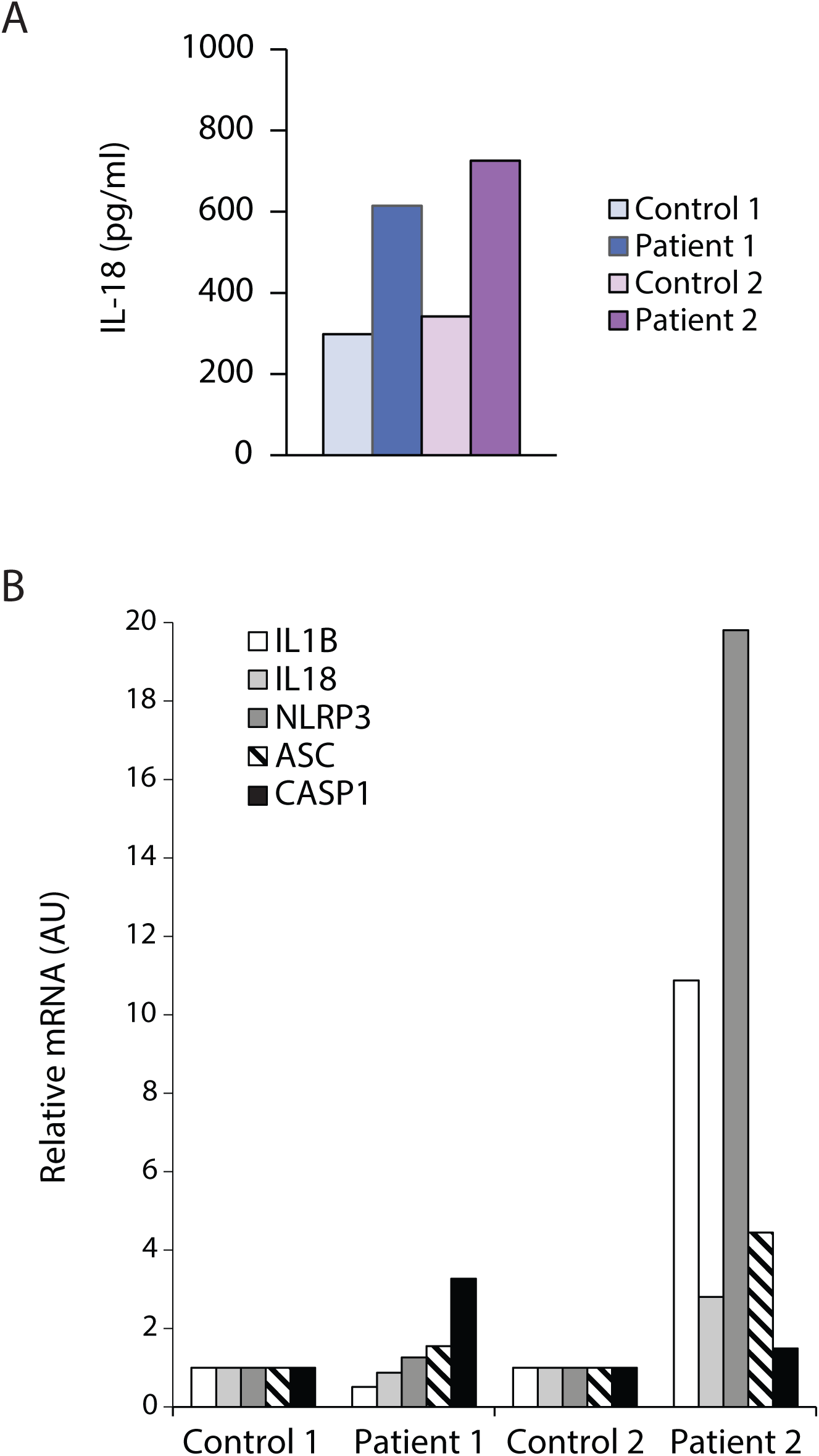
SUPPLEMENTARY FIGURE 3.

## REFERENCES

1. Song HY, Rothe M, Goeddel DV. The tumor necrosis factor-inducible zinc finger protein A20 interacts with TRAF1/TRAF2 and inhibits NF-kappaB activation. Proc Natl Acad Sci U S A1996;93(13):6721-5. [published Online First: 1996/06/25]

2. Heyninck K, Beyaert R. The cytokine-inducible zinc finger protein A20 inhibits IL-1-induced NF-kappaB activation at the level of TRAF6. FEBS Lett 1999;442(2-3):147-50. [published Online First: 1999/02/03]

3. Wertz IE, O’Rourke KM, Zhou H, et al. De-ubiquitination and ubiquitin ligase domains of A20 downregulate NF-kappaB signalling. Nature 2004;430(7000):694–9. doi:10.1038/ nature02794 [published Online First: 2004/07/20]

4. Boone DL, Turer EE, Lee EG, et al. The ubiquitin-modifying enzyme A20 is required for termination of Toll-like receptor responses. Nat Immunol 2004;5(10):1052–60. doi:10.1038/ ni1110 [published Online First: 2004/08/31]

5. Duwel M, Welteke V, Oeckinghaus A, et al. A20 negatively regulates T cell receptor signaling to NF-kappaB by cleaving Malt1 ubiquitin chains. J Immunol 2009;182(12):7718–28. doi:10.4049/jimmunol.0803313 [published Online First: 2009/06/06]

6. Shembade N, Ma A, Harhaj EW. Inhibition of NF-kappaB signaling by A20 through disruption of ubiquitin enzyme complexes. Science 2010;327(5969):1135–9. doi:10.1126/ science.1182364 [published Online First: 2010/02/27]

7. Wertz IE, Newton K, Seshasayee D, et al. Phosphorylation and linear ubiquitin direct A20 inhibition of inflammation. Nature 2015;528(7582):370–5. doi:10.1038/ nature16165 [published Online First: 2015/12/10]

8. Skaug B, Chen J, Du F, et al. Direct, noncatalytic mechanism of IKK inhibition by A20. Mol Cell 2011;44(4):559–71. doi:10.1016/j. molcel.2011.09.015 [published Online First: 2011/11/22]

9. Verhelst K, Carpentier I, Kreike M, et al. A20 inhibits LUBAC-mediated NF-kappaB activation by binding linear polyubiquitin chains via its zinc finger 7. EMBO J 2012;31(19):3845–55. doi:10.1038/emboj.2012.240 [published Online First: 2012/10/04]

10. Krikos A, Laherty CD, Dixit VM. Transcriptional activation of the tumor necrosis factor alpha-inducible zinc finger protein, A20, is mediated by kappa B elements. J Biol Chem 1992;267(25):17971-6. [published Online First: 1992/09/05]

11. Zhou Q, Wang H, Schwartz DM, et al. Loss-of-function mutations in TNFAIP3 leading to A20 haploinsufficiency cause an early- onset autoinflammatory disease. Nat Genet 2016;48(1):67–73. doi:10.1038/ng.3459 [published Online First: 2015/12/08]

12. Lee EG, Boone DL, Chai S, et al. Failure to regulate TNF-induced NF-kappaB and cell death responses in A20-deficient mice. Science 2000;289(5488):2350-4. [published Online First: 2000/09/29]

13. Ma A, Malynn BA. A20: linking a complex regulator of ubiquitylation to immunity and human disease. Nat Rev Immunol 2012;12(11):774–85. doi:10.1038/nri3313[published Online First: 2012/10/13]

14. Das T, Chen Z, Hendriks RW, et al. A20/Tumor Necrosis Factor alpha-Induced Protein 3 in Immune Cells Controls Development of Autoinflammation and Autoimmunity: Lessons from Mouse Models. Front Immunol 2018;9:104. doi:10.3389/fimmu.2018.00104 [published Online First: 2018/03/09]

15. Vande Walle L, Van Opdenbosch N, Jacques P, et al. Negative regulation of the NLRP3 inflammasome by A20 protects against arthritis. Nature 2014;512(7512):69–73. doi:10.1038/nature13322 [published Online First: 2014/07/22]

16. Aeschlimann FA, Batu ED, Canna SW, et al. A20 haploinsufficiency (HA20): clinical phenotypes and disease course of patients with a newly recognised NF-kB-mediated autoinflammatory disease. Ann Rheum Dis 2018;77(5):728–35. doi:10.1136/annrheumdis-2017-212403 [published Online First: 2018/01/11]

17. Kadowaki T, Ohnishi H, Kawamoto N, et al. Haploinsufficiency of A20 causes autoinflam-matory and autoimmune disorders. J Allergy Clin Immunol 2018;141(4):1485–88 e11. doi:10.1016/j.jaci.2017.10.039 [published Online First: 2017/12/16]

18. Takagi M, Ogata S, Ueno H, et al. Haploinsufficiency of TNFAIP3 (A20) by germline mutation is involved in autoimmune lymphoproliferative syndrome. J Allergy Clin Immunol 2017;139(6):1914–22. doi:10.1016/j. jaci.2016.09.038 [published Online First: 2016/11/16]

19. Duncan CJA, Dinnigan E, Theobald R, et al. Early-onset autoimmune disease due to a heterozygous loss-of-function mutation in TNFAIP3 (A20). Ann Rheum Dis 2018;77(5):783–86. doi:10.1136/annrheumdis-2016-210944 [published Online First: 2017/07/01]

20. Ohnishi H, Kawamoto N, Seishima M, et al. A Japanese family case with juvenile onset Behcet’s disease caused by TNFAIP3 mutation. Allergol Int 2017;66(1):146–48. doi:10.1016/j. alit.2016.06.006 [published Online First: 2016/07/28]

21. Shigemura T, Kaneko N, Kobayashi N, et al. Novel heterozygous C243Y A20/TNFAIP3 gene mutation is responsible for chronic inflammation in autosomal-dominant Behcet’s disease. RMD Open 2016;2(1):e000223. doi:10.1136/ rmdopen-2015-000223 [published Online First: 2016/05/14]

22. Trotta L, Hautala T, Hamalainen S, et al. Enrichment of rare variants in population isolates: single AICDA mutation responsible for hyper-IgM syndrome type 2 in Finland. Eur J Hum Genet 2016;24(10):1473–8. doi:10.1038/ejhg.2016.37 [published Online First: 2016/05/05]

23. Trotta L, Martelius T, Siitonen T, et al. ADA2 deficiency: Clonal lymphoproliferation in a subset of patients. J Allergy Clin Immunol 2018;141(4):1534–37 e8. doi:10.1016/j. jaci.2018.01.012 [published Online First: 2018/02/03]

24. Liu X, Salokas K, Tamene F, et al. An AP-MS- and BioID-compatible MAC-tag enables comprehensive mapping of protein interactions and subcellular localizations. Nat Commun 2018;9(1):1188. doi:10.1038/s41467-018- 03523-2 [published Online First: 2018/03/24]

25. Turunen M, Spaeth JM, Keskitalo S, et al. Uterine leiomyoma-linked MED12 mutations disrupt mediator-associated CDK activity. Cell Rep 2014;7(3):654–60. doi:10.1016/j. celrep.2014.03.047 [published Online First: 2014/04/22]

26. Heikkinen T, Kampjarvi K, Keskitalo S, et al. Somatic MED12 Nonsense Mutation Escapes mRNA Decay and Reveals a Motif Required for Nuclear Entry. Hum Mutat 2017;38(3):269–74. doi:10.1002/humu.23157 [published Online First: 2017/01/06]

27. Richards S, Aziz N, Bale S, et al. Standards and guidelines for the interpretation of sequence variants: a joint consensus recommendation of the American College of Medical Genetics and Genomics and the Association for Molecular Pathology. Genet Med 2015;17(5):405–24. doi:10.1038/gim.2015.30 [published Online First: 2015/03/06]

28. Rose PW, Prlic A, Altunkaya A, et al. The RCSB protein data bank: integrative view of protein, gene and 3D structural information. Nucleic Acids Res 2017;5(D1):D271-D81. doi:10.1093/nar/gkw1000 [published Online First: 2016/10/30]

29. Mevissen TET, Kulathu Y, Mulder MPC, et al. Molecular basis of Lys11-polyubiquitin specificity in the deubiquitinase Cezanne. Nature 2016;538(7625):402–05. doi:10.1038/ nature19836 [published Online First: 2016/10/21]

30. Pettersen EF, Goddard TD, Huang CC, et al. UCSF Chimera--a visualization system for exploratory research and analysis. J Comput Chem 2004;25(13):1605–12. doi:10.1002/ jcc.20084 [published Online First: 2004/07/21]

31. He Y, Hara H, Nunez G. Mechanism and Regulation of NLRP3 Inflammasome Activation. Trends Biochem Sci 2016;41(12):1012–21. doi:10.1016/j.tibs.2016.09.002 [published Online First: 2016/09/28]

32. Duong BH, Onizawa M, Oses-Prieto JA, et al. A20 restricts ubiquitination of pro-interleukin-1beta protein complexes and suppresses NLRP3 inflammasome activity. Immunity 2015;42(1):55–67. doi:10.1016/j. immuni.2014.12.031 [published Online First: 2015/01/22]

33. Wang CY, Mayo MW, Korneluk RG, et al. NF-kappaB antiapoptosis: induction of TRAF1 and TRAF2 and c-IAP1 and c-IAP2 to suppress caspase-8 activation. Science 1998;281(5383):1680-3. [published Online First: 1998/09/11]

34. Zarnegar BJ, Wang Y, Mahoney DJ, et al. Noncanonical NF-kappaB activation requires coordinated assembly of a regulatory complex of the adaptors cIAP1, cIAP2, TRAF2 and TRAF3 and the kinase NIK. Nat Immunol 2008;9(12):1371–8. doi:10.1038/ni.1676 [published Online First: 2008/11/11]

35. Hymowitz SG, Wertz IE. A20: from ubiquitin editing to tumour suppression. Nat Rev Cancer 2010;10(5):332–41. doi:10.1038/nrc2775 [published Online First: 2010/04/13]

36. Yamaguchi N, Oyama M, Kozuka-Hata H, et al. Involvement of A20 in the molecular switch that activates the non-canonical NF-small ka, CyrillicB pathway. Sci Rep 2013;3:2568. doi:10.1038/srep02568 [published Online First: 2013/09/07]

37. Yamaguchi N, Yamaguchi N. The seventh zinc finger motif of A20 is required for the suppression of TNF-alpha-induced apoptosis. FEBS Lett 2015;589(12):1369–75. doi:10.1016/j. febslet.2015.04.022 [published Online First: 2015/04/26]

38. Chen X, Arciero CA, Wang C, et al. BRCC36 is essential for ionizing radiation-induced BRCA1 phosphorylation and nuclear foci formation. Cancer Res 2006;66(10):5039–46. doi:10.1158/0008-5472.CAN-05-4194 [published Online First: 2006/05/19]

39. Zhang J, Cao M, Dong J, et al. ABRO1 suppresses tumourigenesis and regulates the DNA damage response by stabilizing p53. Nat Commun 2014;5:5059. doi:10.1038/ ncomms6059 [published Online First: 2014/10/07]

40. Bish RA, Myers MP. Werner helicase-interacting protein 1 binds polyubiquitin via its zinc finger domain. J Biol Chem 2007;282(32):23184–93. doi:10.1074/jbc.M701042200 [published Online First: 2007/06/07]

41. Markkanen E, van Loon B, Ferrari E, et al. Regulation of oxidative DNA damage repair by DNA polymerase lambda and MutYH by cross-talk of phosphorylation and ubiquitination. Proc Natl Acad Sci U S A 2012;109(2):437–42. doi:10.1073/pnas.1110449109 [published Online First: 2011/12/29]

42. Yan K, Li L, Wang X, et al. The deubiquitinating enzyme complex BRISC is required for proper mitotic spindle assembly in mammalian cells. J Cell Biol 2015;210(2):209–24. doi:10.1083/ jcb.201503039 [published Online First: 2015/07/22]

43. Zheng H, Gupta V, Patterson-Fortin J, et al. A BRISC-SHMT complex deubiquitinates IFNAR1 and regulates interferon responses. Cell Rep 2013;5(1):180–93. doi:10.1016/j. celrep.2013.08.025 [published Online First: 2013/10/01]

44. Py BF, Kim MS, Vakifahmetoglu-Norberg H, et al. Deubiquitination of NLRP3 by BRCC3 critically regulates inflammasome activity. Mol Cell 2013;49(2):331–8. doi:10.1016/j. molcel.2012.11.009 [published Online First: 2012/12/19]

45. Jin Z, Li Y, Pitti R, et al. Cullin3-based polyubiquitination and p62-dependent aggregation of caspase-8 mediate extrinsic apoptosis signaling. Cell 2009;137(4):721–35. doi:10.1016/j.cell.2009.03.015 [published Online First: 2009/05/12]

46. Man SM, Kanneganti TD. Converging roles of caspases in inflammasome activation, cell death and innate immunity. Nat Rev Immunol 2016;16(1):7–21. doi:10.1038/nri.2015.7 [published Online First: 2015/12/15]

